# Use of sugarcane mosaic virus for virus-induced gene silencing in maize

**DOI:** 10.1101/2022.02.20.481188

**Authors:** Seung Ho Chung, Shudi Zhang, Hojun Song, Steven A. Whitham, Georg Jander

## Abstract

Previously, sugarcane mosaic virus (SCMV) was developed as a vector for transient expression of heterologous genes in *Zea mays* (maize). Here we show that SCMV can also be applied for virus-induced gene silencing (VIGS) of endogenous maize genes. Comparison of sense and antisense VIGS constructs targeting maize *PDS* (phytoene desaturase) showed that antisense constructs resulted in a greater reduction in gene expression. In a time course of gene expression after infection with VIGS constructs targeting *PDS, Les22* (*Lesion mimic 22*), and *Ij1* (*Iodent japonica1*), efficient expression silencing was observed two, three, and four weeks after infection with SCMV. However, at week five, expression of *Les22* and *Ij1* was no longer significantly reduced compared to control plants. The defense signaling molecule jasmonate-isoleucine (JA-Ile) can be inactivated by 12C-hydroxylation and hydrolysis, and knockout of these genes leads to increased herbivore resistance. JA-Ile hydroxylases and hydrolases have been investigated in Arabidopsis, rice, and *Nicotiana attenuata*. To determine whether the maize homologs of these genes function in plant defense, we silenced expression of *ZmCYP94B1* (predicted JA-Ile hydroxylase) and *ZmJIH1* (predicted JA-Ile hydrolase) by VIGS with SCMV. Although *ZmCYP94B1* and *ZmJIH1gene* expression silencing increased resistance to *Spodoptera frugiperda* (fall armyworm), *Schistocerca americana* (American birdwing grasshopper), and *Rhopalosiphum maidis* (corn leaf aphid), there was no additive effect from silencing the expression of both genes. Further work will be required to determine the more precise functions of these enzymes in regulating maize defenses.

## 1 Introduction

Virus-induced gene silencing (VIGS) is an efficient reverse genetics tool for studying *in vivo* gene function in plants (Hayward et al., 2011). Fragments of a gene of interest are cloned into a virus vector and the endogenous RNA silencing machinery of the host plant causes RNA degradation and reduced expression of the target gene. Since the initial development of tobacco mosaic virus as a vector for gene expression silencing in *Nicotiana benthamiana* (Kumagai et al., 1995), VIGS has been demonstrated using numerous plant-virus combinations (Burch-Smith et al., 2004; Kant and Dasgupta, 2019). Relative to the extensive use of VIGS for studying gene function in dicots, the development of VIGS vectors for *Zea mays* (maize) and other monocot species has been slower. Nevertheless, recent publications describe the use of several viruses for VIGS in maize (Table 1).

**Table 1.** Viruses that have been used to engineer maize VIGS vectors

| **Virus** | **Family** | **References** |
| --- | --- | --- |
| Foxtail mosaic virus | Alphaflexiviridae | Mei et al., 2016 |
| Cucumber mosaic virus | Bromoviridae | Wang et al., 2016 |
| Tobacco rattle virus | Virgaviridae | Zhang et al., 2017 |
| Brome mosaic virus | Bromoviridae | Ding et al., 2018 |
| Barley stripe mosaic virus | Virgaviridae | Jarugula et al., 2018 |
| Maize rayado fino virus | Tymoviridae | Mlotshwa et al., 2020 |
| Sugarcane mosaic virus | Potyviridae | this study |

Sugarcane mosaic virus (SCMV) is a positive-stranded RNA virus in the family Potyviridae that infects sugarcane, maize, and other monocots. An SCMV infectious clone was engineered as a vector for transient gene overexpression in maize (Mei et al., 2019). We have used a version of this vector, SCMV-CS3 (Mohr, 2019; Beernink et al., 2021), which allows cloning of transgenes between the SCMV *P1* and *HC-Pro* cistrons, to enhance maize pest tolerance by overexpressing endogenous maize resistance genes, spider and scorpion toxins, and lectins from other plant species (Chung et al., 2021). Additionally, we used SCMV-VIGS for targeted reduction of gene expression in *Rhopalosiphum maidis* (corn leaf aphids) feeding on maize (Chung and Jander, 2022). However, at that time, we did not investigate whether SCMV-VIGS also can be used to silence gene expression in the host plants.

Degradation of jasmonate-isoleucine (JA-Ile), which functions as an important regulator of plant defenses in many plant species, is a possible VIGS target for increasing plant resistance to insect herbivory. JA-Ile binding to the F-box protein COI1 leads to the degradation of JAZ repressor proteins and induction of defense-related plant gene expression (Howe and Jander, 2008). The level of plant defense induction is regulated by both the biosynthesis and degradation of JA-Ile (Figure 1). Whereas the JAR1 protein conjugates jasmonate and isoleucine to form JA-Ile (Staswick et al., 2002; Koo and Howe, 2009), experiments with *Nicotiana attenuata* showed that JA-Ile can be inactivated by JIH1-mediated cleavage to jasmonate and isoleucine (Woldemariam et al., 2012; Woldemariam et al., 2014). Similarly, *Arabidopsis thaliana* (Arabidopsis) *iar3 ill6* double knockouts, which are defective in JA-Ile deconjugation, exhibited increased JA-Ile accumulation (Marquis et al., 2020). The *Oryza sativa* (rice) genes *IAR3* and *AH8* encode similar JA-Ile hydrolases (Hazman et al., 2019).

**Figure 1.**
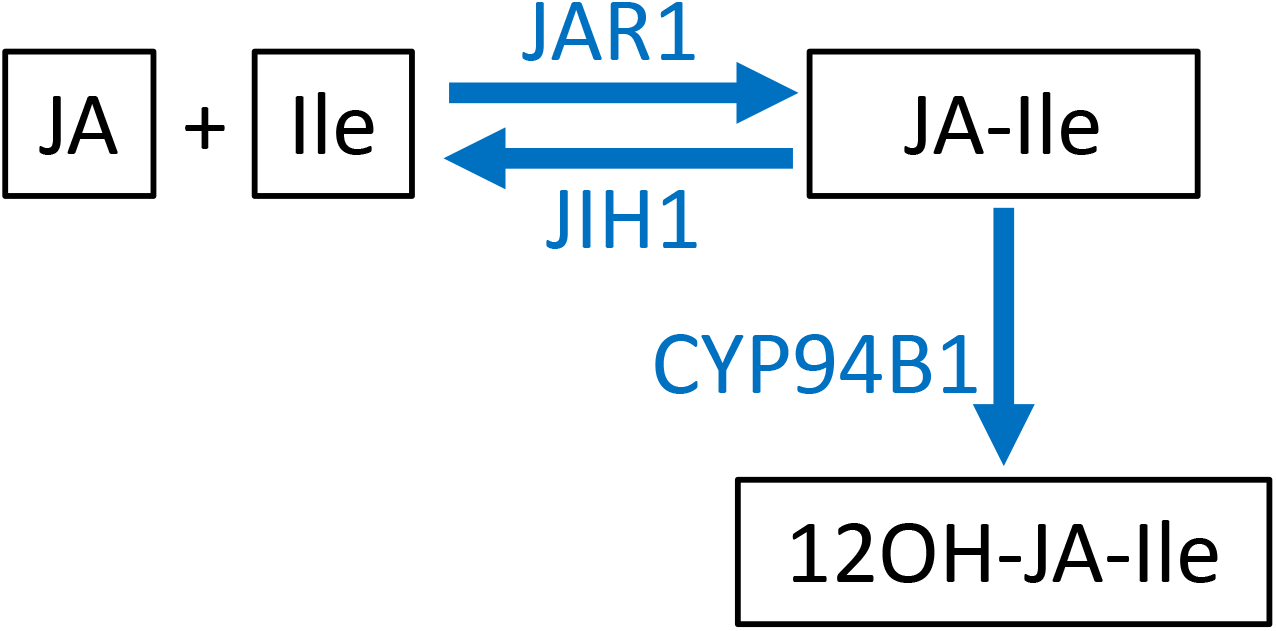
Jasmonate-isoleucine (JA-Ile) synthesis and degradation. Abundance of the plant defense signaling molecule JA-Ile is affected by both biosynthesis and degradation. Whereas JAR1 conjugates jasmonate (JA) and isoleucine (Ile) to form JA-Ile, JIH1 catalyzes the reverse reaction. CYP94B1 inactivates JA-Ile by oxidation to form 12-hydroxy-JA-Ile (12OH-JA-Ile).

Research with Arabidopsis showed that JA-Ile can be hydroxylated by CYP94B1 and CYP94B3 to produce 12-hydroxy jasmonate-isoleucine (12OH-JA-Ile), which is less efficient than JA-Ile in eliciting COI1-mediated JAZ protein degradation (Kitaoka et al., 2011; Koo et al., 2011; Koo et al., 2014; Marquis et al., 2020). A rice gene, CYP94B5 also encodes a JA-Ile C12-hydroxylase (Hazman et al., 2019). In maize, a dominant *CYP94B1* mutation, known as *Tasselseed5 (Ts5*), causes increased gene expression, lower JA-lle, and higher 12OH-JA-lle accumulation than in wildtype plants (Lunde et al., 2019).

Higher pest tolerance in plants can be achieved by increasing the biosynthesis or decreasing the inactivation of JA-Ile. Expression of *JAR1a* and *JAR1b*, two of the five maize genes predicted to encode JA-Ile conjugating enzymes (Borrego and Kolomiets, 2016), was strongly induced by caterpillar feeding (Tzin et al., 2017). Transient overexpression of these genes in maize using SCMV caused reduced growth of *Spodoptera frugiperda* (fall armyworm) larvae (Chung et al., 2021). Conversely, RNA interference targeting *JIH1* in *N. attenuata* made these plants more resistant to both *Manduca sexta* (tobacco hornworm) and *Spodoptera littoralis* (Egyptian cotton leafworms) (Woldemariam et al., 2012), and silencing expression of *N. attenuata* JA hydroxylases increased resistance to *Spodoptera litura* (Tang et al., 2020). Similarly, knockout of the two Arabidopsis JA-Ile hydrolase genes, *IAR3* and *ILL6*, decreased *S. littoralis* caterpillar growth (Marquis et al., 2020) and knockout of jasmonate hydroxylases increased resistance to multiple biotic and abiotic stresses (Caarls et al., 2017; Smirnova et al., 2017; Marquis et al., 2021).

Here we describe the development of an SCMV-VIGS protocol for maize and demonstrate its research utility by silencing the expression of two JA-Ile inactivating genes, the predicted maize homologs of *JIH1* and *CYP94B1*. Reduced expression of *ZmJIH1* and *ZmCYP94B1* caused higher expression of known jasmonate-regulated defense genes and elevated resistance to feeding by species in three different insect orders: *S. frugiperda* (Lepidoptera), *Schistocerca americana* (American birdwing grasshopper, Orthoptera), and *R. maidis* (Hemiptera) (Figure 2).

**Figure 2.**
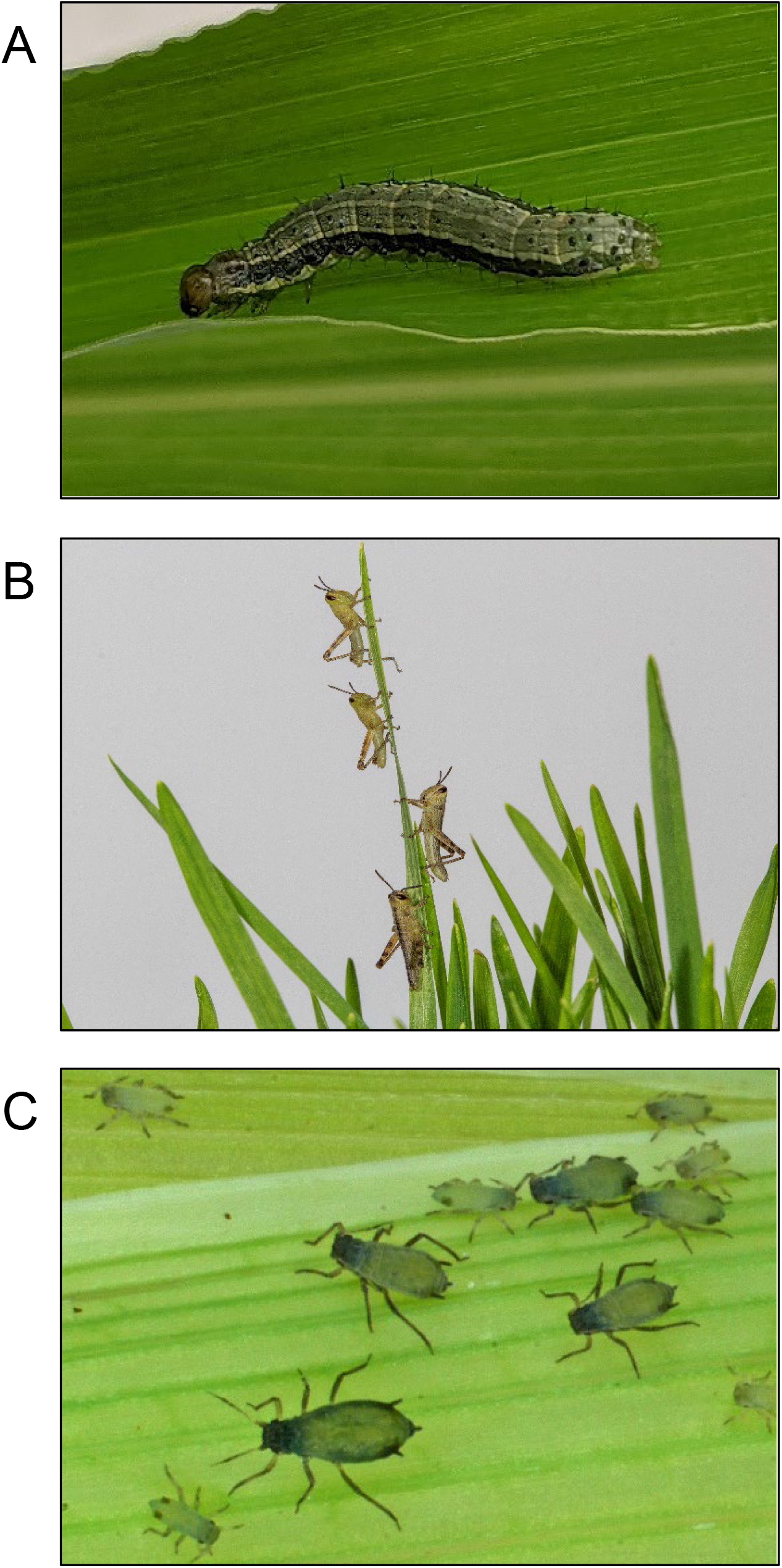
Insects used in this study. (A) *Spodoptera frugiperda* (fall armyworm) caterpillars (credit: Seung Ho Chung), (B) *Schistocerca americana* (American birdwing grasshopper) nymphs (photo credit: Brandon Woo), (C) mixed-instar *Rhopalosiphum maidis* (corn leaf aphids; photo credit: Meena Haribal).

## 2 Materials and Methods

### 2.1 Plants, insects, and growth conditions

Maize (*Zea mays*) inbred line P39 for SCMV infection and VIGS experiments was grown in a soil mix that was prepared in batches consisting of: 0.16 m^3^ Metro-Mix 360 (Scotts, www.scotts.com), 0.45 kg finely ground lime, 0.45 kg Peters Unimix (Griffin Greenhouse Supplies, www.griffins.com), 68 kg Turface MVP (Banfield-Baker Corp., www.banfieldbaker.com), 23 kg coarse quartz sand, and 0.018 m^3^ pasteurized field soil. Plants were maintained in a growth chamber at 23 °C with a 16:8 h light:dark cycle.

*Spodoptera frugiperda* eggs were purchased from Benzon Research (www.benzonresearch.com) and were placed on artificial diet (Fall Armyworm Diet, www.southlandproducts.net) in an incubator at 28°C for hatching. A colony of a genome-sequenced *R. maidis* lineage (Chen et al., 2019) was maintained on inbred line P39 or sweet corn variety Golden Bantam (Burpee Seeds, www.burpee.com) at 23°C under 16:8 h light:dark cycle. A colony of *S. americana*, started with insects originally collected from St. Augustine, Florida (29°39′30.4″N 81°17′16.0″W and 29°40′16.3″N 81°15′37.0″W) in October 2018 and was maintained on wheat grass, Romaine lettuce, and wheat bran at 30°C at 12:12 hr light-dark cycle at the USDA-approved quarantine facility in the Department of Entomology at Texas A&M University. The grasshopper egg pods were transported to Boyce Thompson Institute under the USDA-APHIS permit P526P-21-06015, and the hatchlings were used for experiments.

### 2.2 Maize infection with VIGS constructs

Fragments of the genes to be silenced, *PDS* (GRMZM2G410515), *Les22* (GRMZM2G044074), *Ij* (GRMZM2G004583), *JIH1* (GRMZM2G090779) and *CYP94B1* (GRMZM2G177668) (Table S1), were identified using pssRNAit (www.zhaolab.org/pssRNAit/ Ahmed et al., 2020). Gene fragments for VIGS were chosen such that they have no significant off-target matches elsewhere in the maize genome. In the case of antisense constructs, gene fragments were chosen such that they both conserve the open reading frame when cloned in SCMV and have no in-frame stop codons. cDNA of maize inbred line B73 was amplified using the primers listed in Table S2. A 363 bp fragment for simultaneous silencing of *CYP94B1* and *JIH1* (Table S2) was synthesized by Twist Bioscience (www.twistbioscience.com). The pSCMV-CS3 vector (Chung et al., 2021), which expresses full-length SCMV RNA from the cauliflower mosaic virus 35S promoter, was cut with the restriction enzymes *Psp*OMI and *PmeI* (New England Biolabs, www.neb.com), the amplified gene fragments were cloned into the cut site such that they were in-frame with the viral RNA, and the constructs were transformed into *Escherichia coli* Top10 competent cells (www.thermofisher.com). A fragment of *GFP* (Table S2) was cloned into SCMV-CS3 as a control

For biolistic plant transformation (as described by Chung et al., 2021), plasmid DNA carrying the SCMV constructs was coated onto 3 mg 1.0 μm diameter gold particles. The gold particles were distributed onto five particle bombardment macrocarriers and allowed to air dry. One-week-old P39 seedlings were placed into a Biolistic PDS-1000/He system (www.biorad.com), randomly oriented so that the adaxial or abaxial surface faced upward for bombardment (Figure 3A). Macrocarriers and 1,100 psi rupture disks were placed into the biolistic system and leaves were bombarded at a distance of 6 cm between the stopping screen and the leaves.

**Figure 3.**
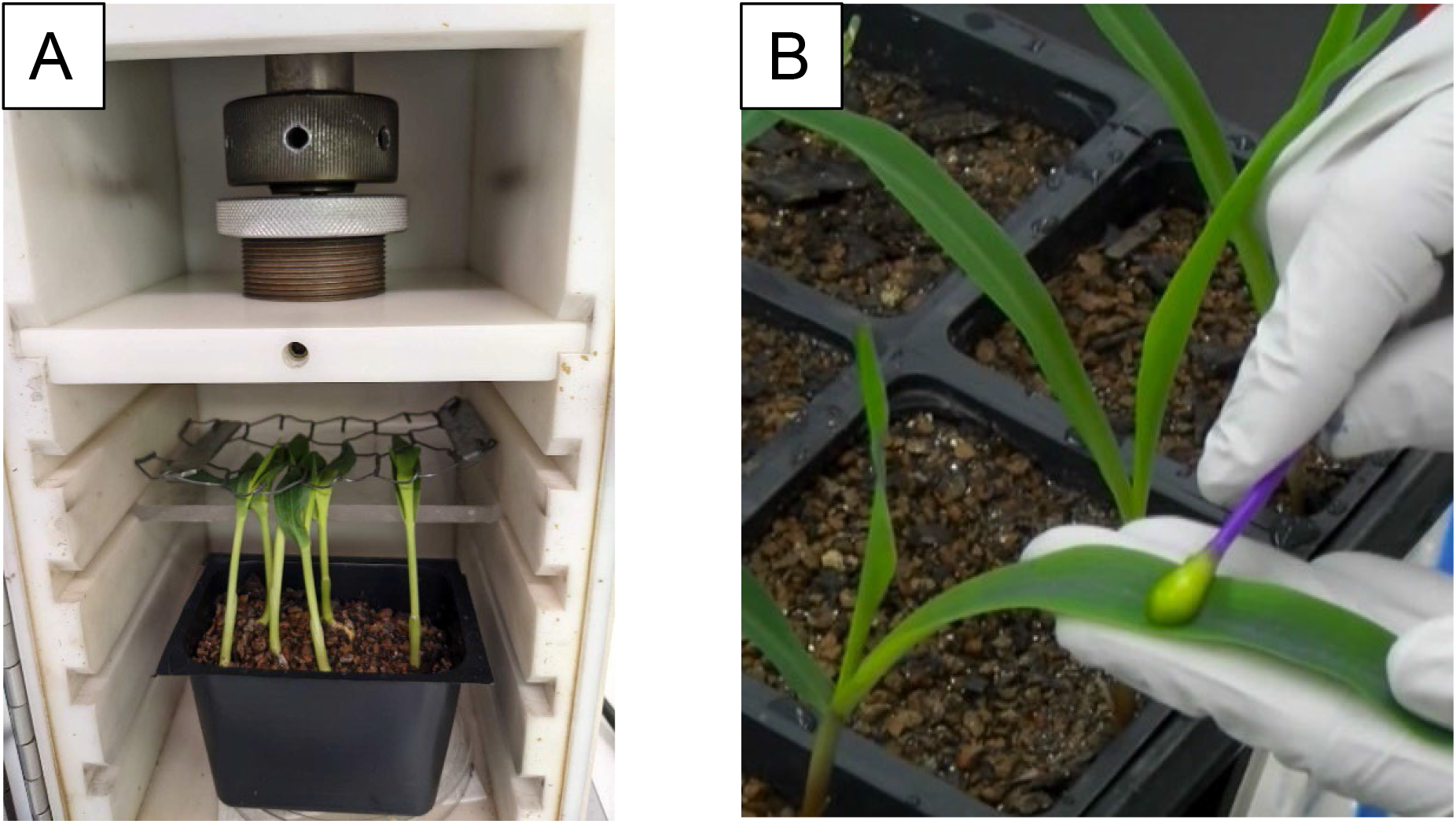
Infection of maize with sugarcane mosaic virus (SCMV). (A) For initial infection, plasmid DNA encoding SCMV constructs was transformed into one-week-old maize seedlings using a Biolistic PDS-1000/He system; (B) To generate plants for experimental assays, leaves of maize seedlings were dusted with carborundum and a cotton swab was used to rub on sap from SCMV-infected maize leaves.

For further propagation and insect experiments, sap of SCMV-infected maize plants was prepared by grinding 0.5 g leaf tissue in 5 ml of 50 mM pH 7.0 potassium phosphate buffer. One-week-old P39 maize plants were mechanically infected by dusting the leaves with 600-mesh carborundum and rubbing the SCMV-containing plant sap onto the surface with a cotton swab (Figure 3B). Successfully infected plants were identified by the development of viral symptoms three weeks later.

### 2.3 Analysis of gene expression by quantitative RT-PCR

Two, three, four, and five weeks after SCMV infection by rub-inoculation, the seventh or eighth leaves of infected plants were collected, flash-frozen in liquid nitrogen, and stored at −80 °C. For induction experiments, two 5-day-old *S. frugiperda* caterpillars were added in clip cages (2.5 x 3.0 cm) that were placed on the seventh or eighth leaves of infected plants three weeks after SCMV infection. Control treatments without herbivory received empty cages. Caterpillars were removed 24 hours later and about 100 mg of damaged tissue was harvested from each plant. RNA was extracted using TRIzol Reagent (www.invitrogen.com) and treated with RQ1 RNase-free DNase (www.promega.com). One microgram of RNA was used to synthesize first-strand cDNA using High-Capacity cDNA Reverse Transcription Kit (Applied Biosystems; www.thermofisher.com) with random primers. Primers used to measure the expression of the target genes are listed in Table S2. The reactions consisted of 5.0 μl of the PowerUp SYBR Green PCR master mix (Applied Biosystems), 0.6 μl primer mix (300 nM for the final concentration of each primer) and 2 μl of cDNA (1:10 dilution with nuclease-free H2O) in 10 μl total volume. Template-free reactions were included as negative controls. The PCR amplification was performed on QuantStudio 6 Flex Real-Time PCR Systems (Applied Biosystems) with the following conditions: 2 min at 50°C, 2 min at 95°C, 40 cycles of 95°C for 15 sec, and 60°C for 1 min. Primer specificity was confirmed by melting curve analysis. Mean cycle threshold values of duplicates of each sample were normalized using two reference genes, *Actin* and *EF1-α*. Relative gene expression values were calculated using 2^-ΔΔCt^ method (Livak and Schmittgen, 2001).

### 2.4 Insect bioassays

Four-week-old maize plants, three weeks after infection with SCMV, were used for insect bioassays. For *S. frugiperda* growth assays, each maize plant received five 2-day-old caterpillars and was enclosed in a perforated plastic bag (13 cm x 61 cm; www.clearbags.com). After one week of feeding, the fresh mass of the surviving caterpillars was measured and the average mass of caterpillars from each plant was used as a biological replicate in statistical comparisons of maize plants infected with different SCMV VIGS constructs. For *S. americana* assays, five 1 to 3-day old nymphs were weighed and placed onto maize plants that were covered with perforated plastic bags. The *S. americana* nymphs were weighed again 7 days later. The average weight of nymphs in each bag at the beginning and end of the experiment was used as a biological replicate to calculate the relative growth rate (RGR). RGR of grasshoppers was calculated according to the formula: RGR = (ln W2 - ln W1)/(t2 - t1), where W1 and W2 are the average insect wet weights on each plant at times t1 and t2. For aphid bioassays, eight 10-day-old apterous adult *R. maidis* were placed on each SCMV-infected plant and enclosed using perforated plastic bags. After one week, the total number of aphids on each maize plant was counted.

### 2.5 Phylogenetic tree construction

Protein sequences of known JA-Ile hydrolases were downloaded from www.Arabidopsis.org (Arabidopsis;), www.rice.uga.edu (rice;), and www.ncbi.nlm.nih.gov/genbank/ (*Nicotiana attenuata;*). Based on BLAST searches with *N. attenuata* JIH1, sequences of the five most similar maize inbred line B73 proteins (GRMZM2G090779, GRMZM5G833406, GRMZM2G091540, GRMZM2G476538, and GRMZM2G125552) were downloaded from www.maizegdb.org. MEGA11 (Tamura et al., 2021; www.megasoftware.net) was used to construct a phylogeny using the Maximum Likelihood method and the Whelan And Goldman model (Whelan and Goldman, 2001). A discrete Gamma distribution was used to model evolutionary rate differences among sites (5 categories (+G, parameter = 0.9777)). All positions with less than 95% site coverage were eliminated from the analysis, leaving a total of 394 positions in the final dataset. The bootstrap consensus values (Felsenstein, 1985) are percentages inferred from 1000 replicates.

### 2.6 Statistical Analysis

Raw data underlying the bar graphs in Figures 4, 5, 6, 8, and 9 are in Tables S3-S7. All statistical analyses were conducted using R (R Core Team, 2017). Gene expression data were log2 transformed before the statistical analysis, but untransformed data are presented in the figures. Data for gene expression and insect bioassay were analyzed using *t*-tests or analysis of variance (ANOVA) followed by Tukey’s test.

### 2.7 Accession Numbers

Sequences of maize genes and proteins were downloaded from maizeGDB (www.maizeGDB.org) and include GRMZM2G410515 (*PDS*), GRMZM2G044074 (*Les22*), GRMZM2G004583 (*Ij*), GRMZM2G090779 (*JIH1*), GRMZM2G177668 (*CYP94B1*), GRMZM2G090779 (*JIH1*), GRMZM5G833406, GRMZM2G091540, GRMZM2G476538, and GRMZM2G125552.

### 2.8 List of Supplementary Materials

Table S1. Sequences of inserts in VIGS constructs

Table S2. Primers used for cloning and qRT-PCR

Table S3. Raw data underlying the graphs in Figure 4

Table S4. Raw data underlying the graphs in Figure 5

Table S5. Raw data underlying the graphs in Figure 6

Table S6. Raw data underlying the graphs in Figure 8

Table S7. Raw data underlying the graphs in Figure 9

## 3 Results and Discussion

We used the previously described SCMV-CS3 vector (Mohr, 2019; Beernink et al., 2021; Chung et al., 2021) to determine whether SCMV can be used for gene expression silencing in maize. As SCMV is translated as a single polycistronic protein prior to being cleaved by virus-encoded proteases (Mei et al., 2019), sense gene fragments must be cloned such that they are in-frame with the viral coding sequence. Antisense gene fragments for SCMV VIGS must be chosen carefully, so that the resulting coding sequence is in frame with the virus proteins and there are no in-frame stop codons on the otherwise non-coding strand of the gene. Following these criteria, we cloned sense and antisense *PDS* (phytoene desaturase; GRMZM2G410515) gene fragments (Table S1) between the *P1* and *HC-Pro* cistrons in SCMV-CS3 (Figure 4A). *PDS* gene expression was measured by quantitative reverse transcriptase polymerase chain rection (qRT-PCR) two weeks (Figure 4B) and three weeks (Figure 4C) after infection. Although both constructs reduced maize *PDS* gene expression relative to maize infected with an antisense *GFP* control, gene expression was significantly lower when using the antisense *PDS* construct. Therefore, all subsequent experiments were conducted with target gene fragments cloned into the antisense orientation.

**Figure 4.**
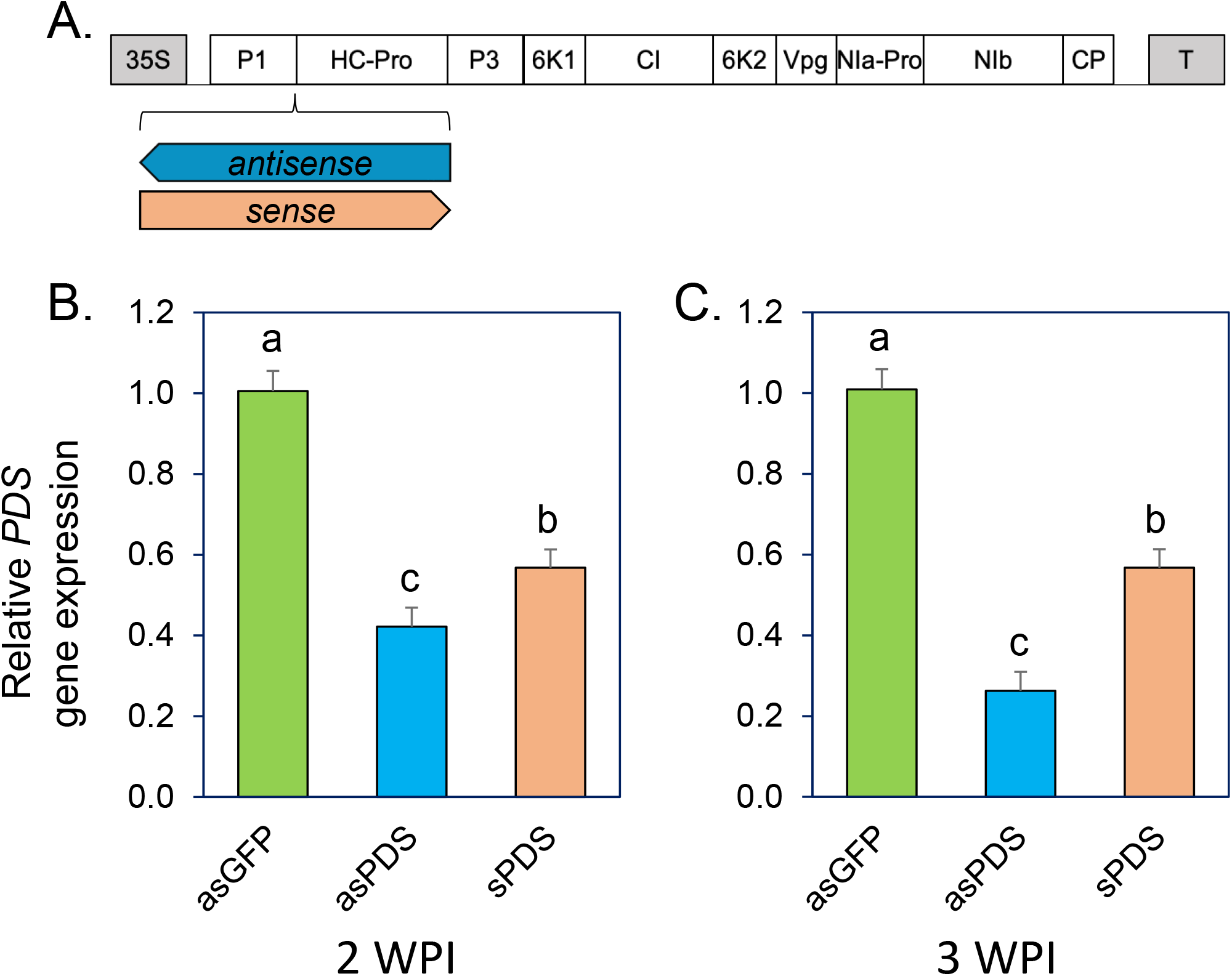
Antisense sugarcane mosaic virus (SCMV) constructs silence gene expression more efficiently. (A) The pSCMV-CS3 binary vector encodes SCMV expressed from the cauliflower mosaic virus 35S promoter. Constructs targeting *PDS* were cloned between the SCMV P1 and HC-Pro cistrons in the antisense (asPDS) and sense (sPDS) orientation and were used to infect maize inbred line P39. Control maize plants were infected with SCMV carrying a similarly-sized GFP fragment in the antisense orientation. Expression of maize PDS in infected leaves was measured at two (B) and three (C) weeks post-infection (WPI). Means +/- s.e. of n = 6; different letters above the bars indicate significant differences, ANOVA followed by Tukey’s HSD test.

To determine the length of time over which effective expression silencing is observed, *PDS*, *Les22* (*lesion mimic 22*, uroporphyrinogen decarboxylase, GRMZM2G044074), and *Ij1* (*Iodent japonica1*, GRMZM2G004583) were targeted with antisense VIGS constructs. At two, three and four weeks after infection, there was a 50% to 70% reduction in the expression of the three tested genes relative to control plants infected with an SCMV-*GFP* (Figure 5A-C). However, five weeks after virus infection, only *PDS* expression was significantly reduced relative to the controls. Subsequent experiments were done with maize plants three weeks after inoculation with SCMV VIGS constructs.

**Figure 5.**
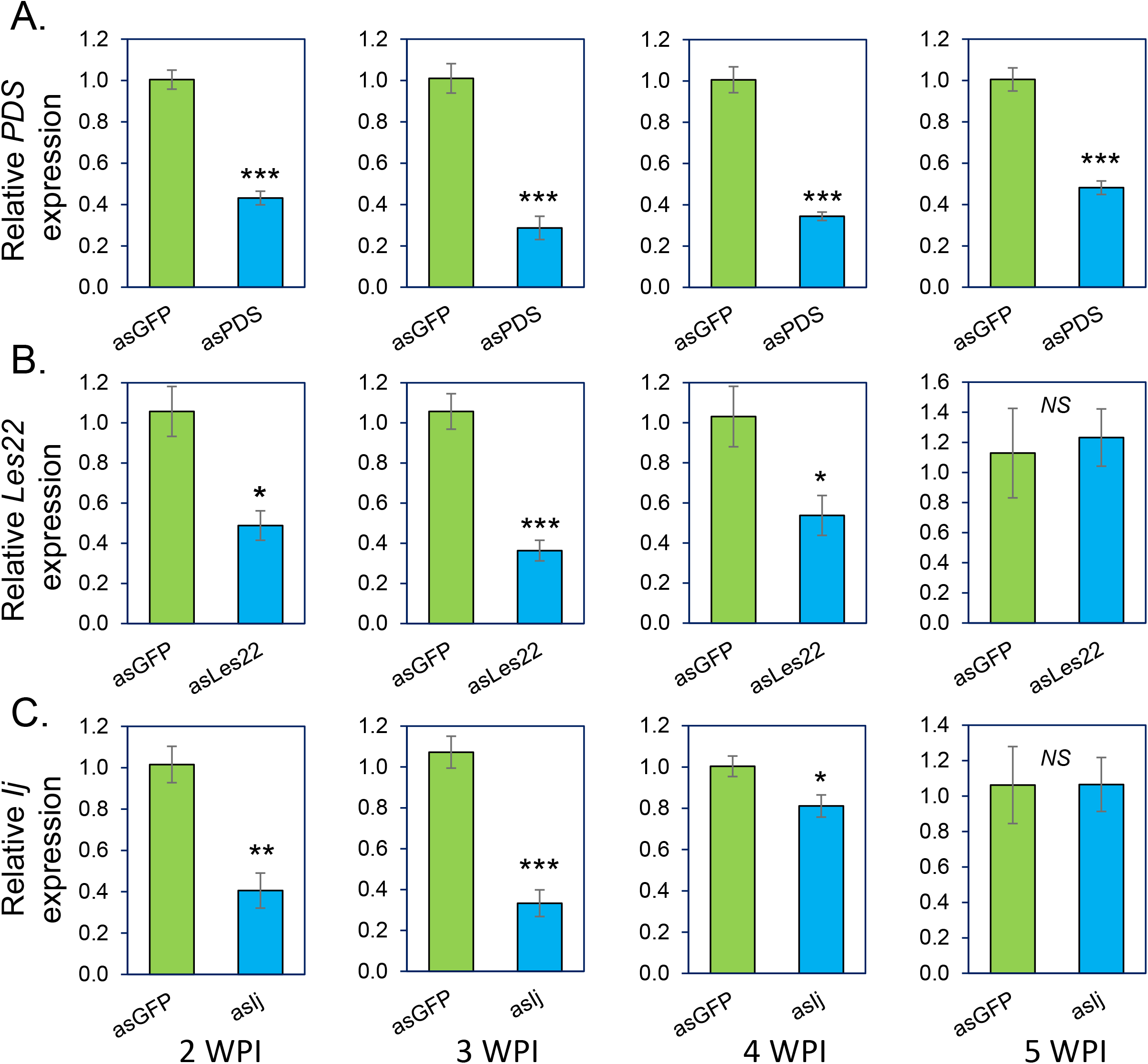
Virus induced gene silencing (VIGS) in maize using sugarcane mosaic virus (SCMV). Maize inbred line P39 was rub-inoculated with SCMV carrying fragments of (A) *PDS*, (B) *Les22*, or (C) *Ij*, and gene expression was measured at two, three, four, and five weeks post-infection (WPI). Control plants were infected with SCMV carrying a fragment of GFP. Mean +/- s.e. of n = 4-5; *: P<0.05, **: P<0.01, ***: P<0.001, and *NS.:* non-significant, as determined by *t*-tests.

*Ts5*, a dominant mutation that increases maize 12OH-JA-Ile accumulation (Lunde et al., 2019), was identified as a homolog of the Arabidopsis JA-Ile 12C-hydroxylase *CYP94B1* (Koo et al., 2014). We cloned a fragment of this gene (*ZmCYP94B1;* GRMZM2G177668) into the SCMV-CS3 vector in the antisense orientation for VIGS experiments. *ZmCYP94B1* expression was significantly reduced by VIGS, with and without feeding by *S. frugiperda* (Figure 6A). Expression of the maize defensive genes, *MPI (Maize Proteinase Inhibitor*, GRMZM2G028393; Cordero et al., 1994; Tamayo et al., 2000; Shivaji et al., 2010) and *RIP2 (Ribosome Inactivating Protein2*, GRMZM2G119705; Chuang et al., 2014) is upregulated in response to JA treatment and insect feeding. Both genes were expressed at a higher level in *ZmCYP94B1-silenced* plants in response to *S. frugiperda* feeding than in the corresponding SCMV-GFP controls (Figure 6B,C), indicating that defense induction is enhanced. *Spodoptera frugiperda* (Figure 6D), *S. americana* (Figure 6E), and *R. maidis* (Figure 6F) grew less well on *ZmCYP94B1-silenced* maize than on plants infected with SCMV-GFP, consistent with previous reports showing that MPI and RIP2 inhibit insect growth (Vila et al., 2005; Chuang et al., 2014; Quilis et al., 2014; Chung et al., 2021). However, it is likely that, in addition to these two proteins, other jasmonate-regulated maize defenses are increased in response to *ZmCYP94B1* expression silencing.

**Figure 6.**
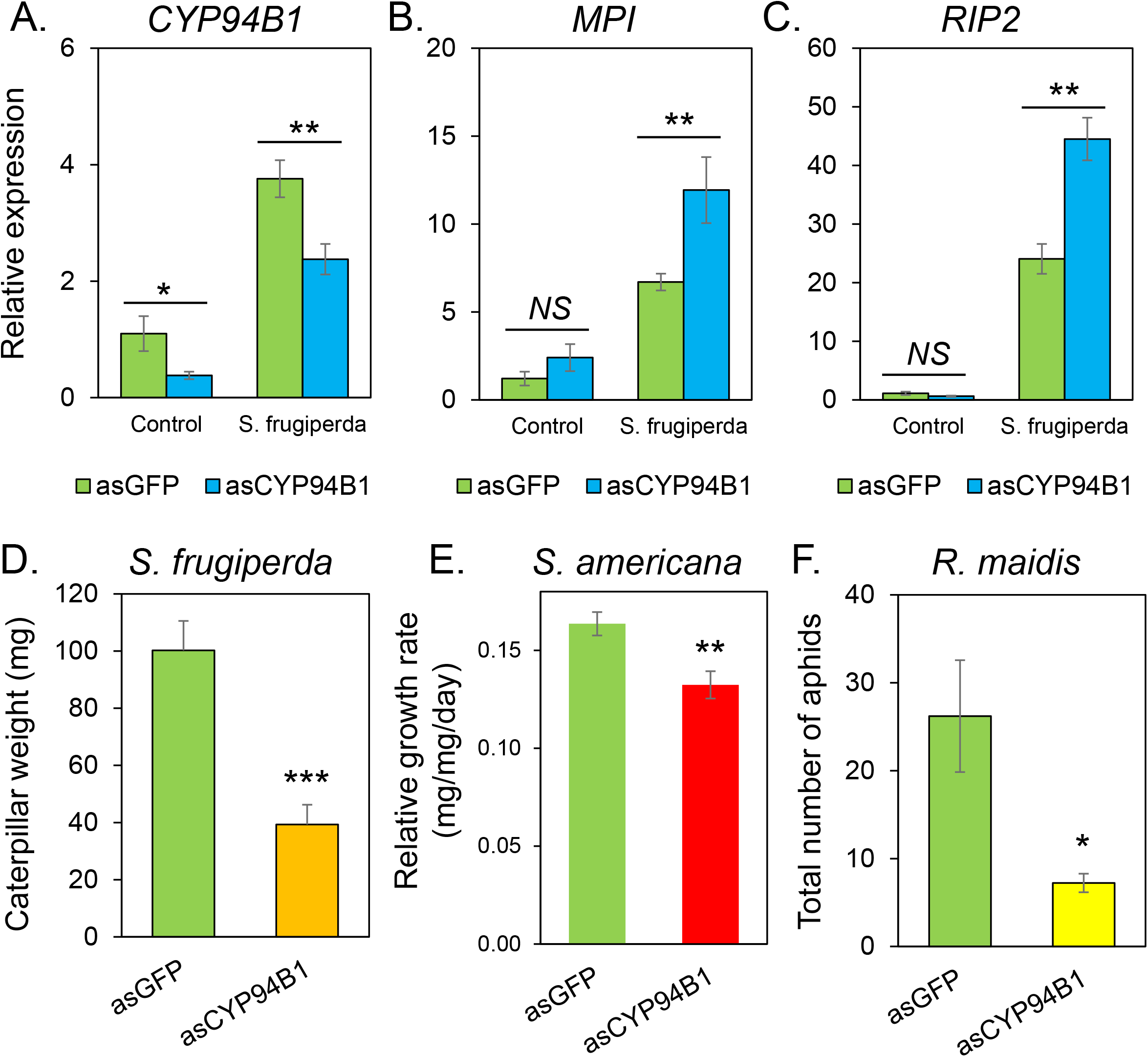
VIGS of *CYP94B1* and its effects on defense gene expression and insect growth. Maize inbred line P39 was rub-inoculated with SCMV carrying a *CYP94B1* fragment, or *GFP* as a control, in the antisense orientation. Expression of (A) *CYP94B1*, (B) *MPI*, and (C) *RIP2* was measured by qRT-PCR three weeks after SCMV infection. Mean +/- s.e. of N = 5. (D) *Spodoptera frugiperda* caterpillar weight after one week, mean +/- s.e. of N = 9. (E) Relative growth rate of *Schistocerca americana* nymphs, mean +/- s.e. of N = 10 (control) and 12 (*asCYP94B1*) (F) Total number of aphids one week after aphids were place on plants, mean +/- s.e. of N = 10 (control) and N = 9 (*asCYP94B1*); the control sample is the same as in Figure 8F; *: P<0.05, **: P<0.01, ***: P<0.001, and *NS.:* non-significant, as determined by *t-*tests.

Our observation of decreased *S. frugiperda* growth due to *ZmCYP94B1* expression silencing is different from what was observed with Arabidopsis, where *S. littoralis* larval growth on a *cyp94B1 cyp94B3 cyp94C1* triple mutant was improved relative to wildtype plants (Marquis et al., 2020). The faster caterpillar weight gain on this triple mutant was ascribed to the higher expression of *JAZ* repressor proteins, which negatively regulate defense-induced genes, in the triple mutant. Further experiments will need to be done to determine whether or not maize *JAZ* gene expression is similarly upregulated when *ZmCYP94B1* expression is reduced. However, the observation of decreased caterpillar growth suggests that this aspect of maize defense regulation is different from that which has been found in Arabidopsis.

A phylogenetic analysis (Figure 7) showed that *ZmJIH1* (GRMZM2G090779) is the closest maize homolog of *Nicotiana attenuata JIH1* (Woldemariam et al., 2012), rice *AH8* (Hazman et al., 2019), and Arabidopsis *IAR3* (Marquis et al., 2020), which have previously been identified as JA-Ile hydrolases. We cloned a fragment of this gene into SCMV-CS3 for VIGS experiments. Relative to plants infected with SCMV-*GFP*, those infected with *SCMV-JIH1*, had a 50% reduction in *ZmJIH1* expression in the absence of *S. frugiperda* feeding (Figure 8A). Both *MPI* and *RIP2* were expressed at a higher level in *ZmJIH1-*silenced plants in response to *S. frugiperda* feeding than in the corresponding SCMV-GFP controls (Figure 8B,C). The three tested insect species, *S. frugiperda*, *S. americana*, and *R. maidis* all grew less well on *ZmJIH1-* silenced plants than on corresponding control plants (Figure 8D-F). Thus, similar to what has been observed in other plant species (Woldemariam et al., 2012; Marquis et al., 2020), maize *Zm*JIH1 is a negative regulator jasmonate-induced gene expression and insect resistance.

**Figure 7.**
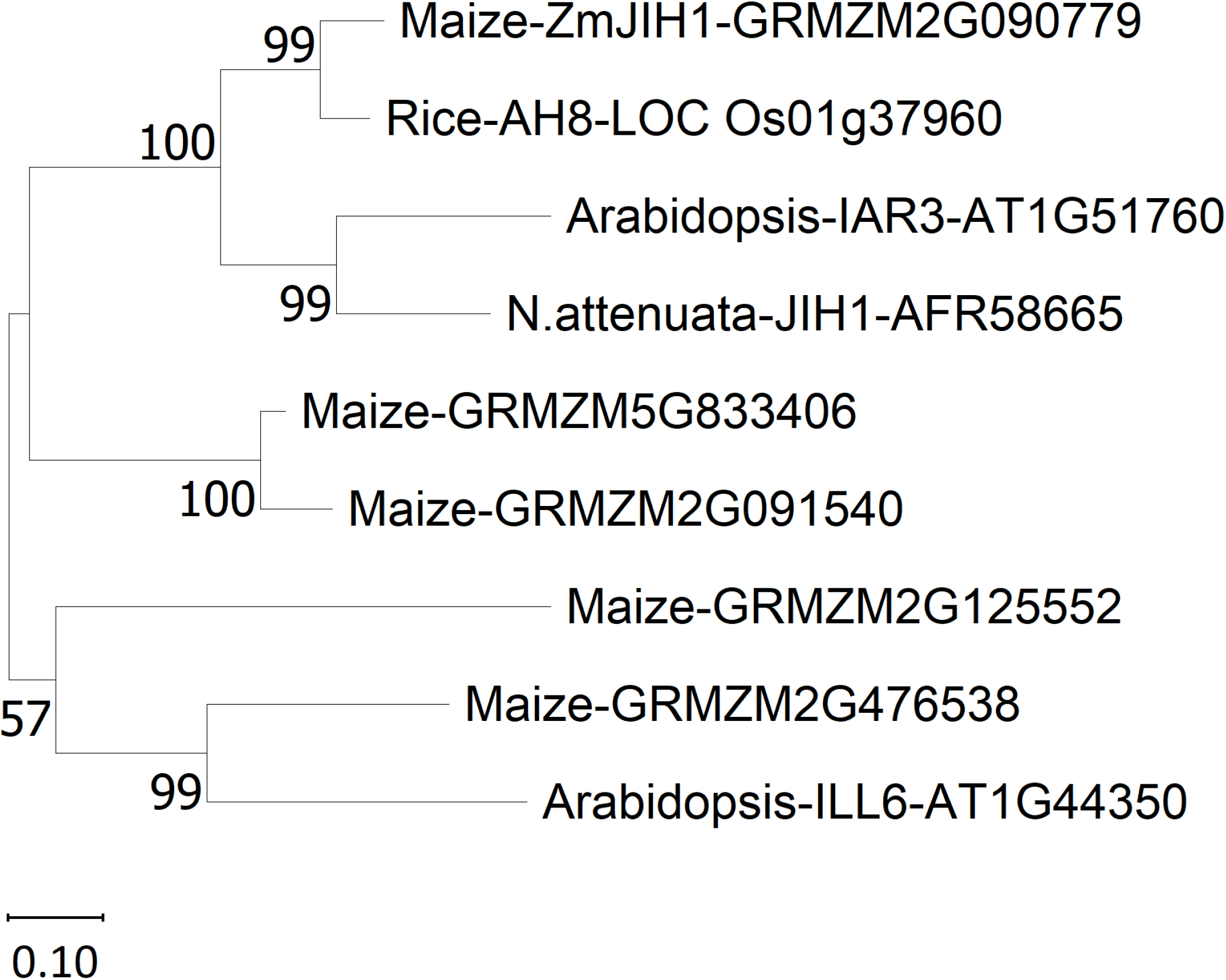
Maximum likelihood tree for identifying a maize JA-Ile hydrolase candidate. Arabidopsis, rice, and *N. attenuata* proteins with known JA-Ile hydrolase activity were in BLAST searches to identify the most similar maize proteins. Based on an unrooted phylogeny constructed with MEGA 11 (www.megasoftware.net), GRMZM2G090779 (*ZmJIH1*) was chosen as the most likely maize protein to have JA-Ile hydrolase activity. The bootstrap consensus values are percentages inferred from 1000 replicates.

**Figure 8.**
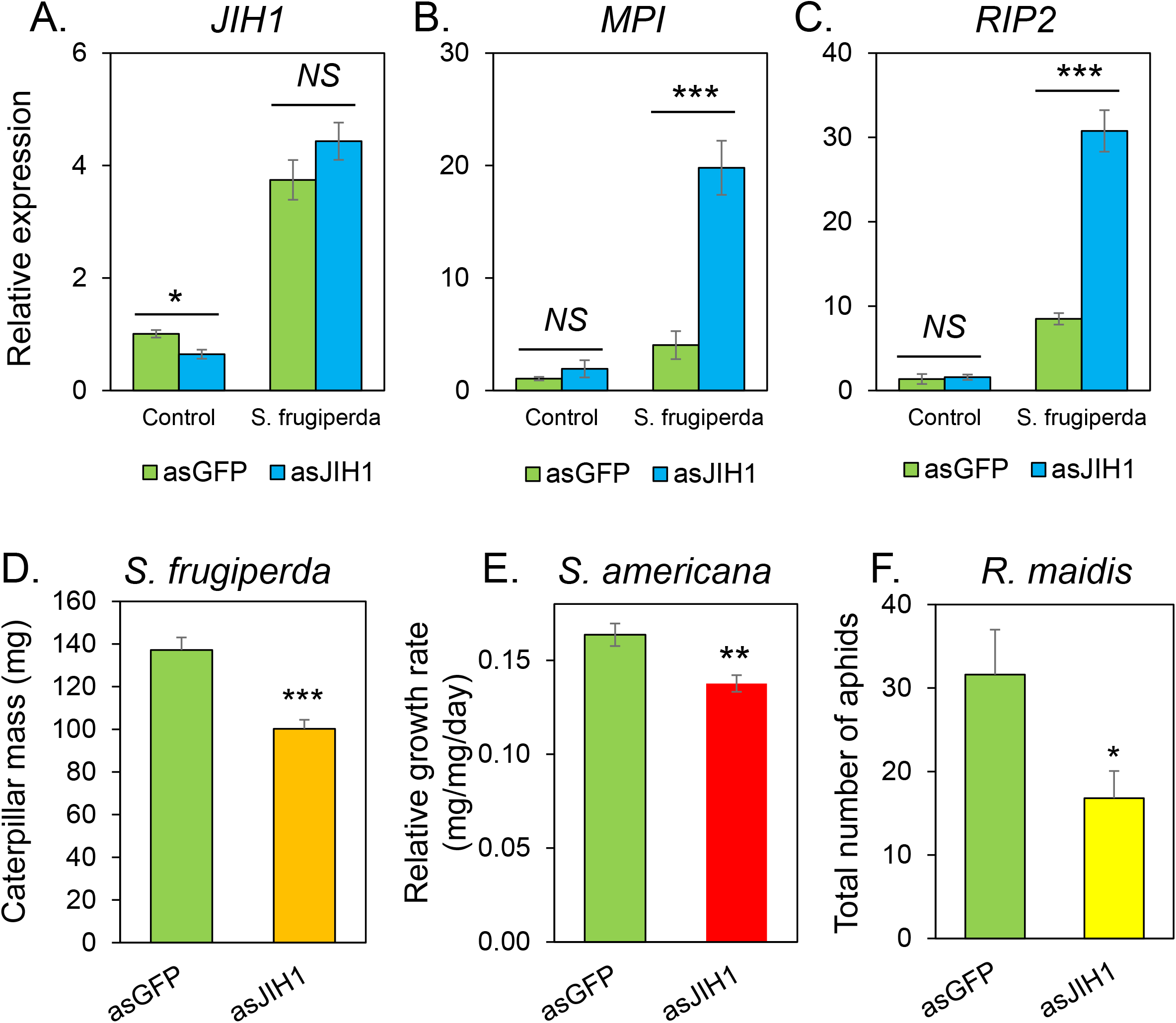
VIGS of *JIH1* and its effects on defense gene expression and insect growth. Maize inbred line P39 was rub-inoculated with SCMV carrying a *JIH1* fragment, or *GFP* as a control, in the antisense orientation. Expression of (A) *JIH1*, (B) *MPI*, and (C) *RIP2* was measured by qRT-PCR three weeks after SCMV infection. Mean +/- s.e. of N = 5. (D) *Spodoptera frugiperda* caterpillar weight after one week, mean +/- s.e. of N = 14 (control) and N = 17 (*JIH1;*). (E) Relative growth rate of *Schistocerca americana* nymphs, mean +/- s.e. of N = 8 (control) and 15 (*JIHΓ*) (F) Total number of aphids one week after aphids were place on plants, mean +/- s.e. of N = 12; the control sample is the same as in Figure 6F; *: P<0.05, **: P<0.01, ***: P<0.001, and *NS.:* nonsignificant, as determined by *t*-tests.

To determine whether simultaneous silencing of *ZmCYP94B1* and *ZmJIH1* expression has an additive effect on insect resistance, we made a VIGS construct targeting both of these genes. The two-gene construct silenced both genes as effectively as each of the single-gene VIGS constructs, SCMV-*CYP94B1* (Figure 9A) and SCMV-*JIH1* (Figure 9B). However, there was no additive effect on insect resistance. The reduction in *S. frugiperda* caterpillar growth (Figure 9C) and *R. maidis* reproduction (Figure 9D) did not differ between the single-gene and two-gene VIGS constructs. By contrast, in Arabidopsis, where knockout of JA-Ile hydrolase and JA-Ile hydroxylase activity had opposite effects on *S. littoralis* larval growth, knockout of both enzymatic activities resulted in plants that were not significantly different from wildtype in their caterpillar resistance (Marquis et al., 2020).

**Figure 9.**
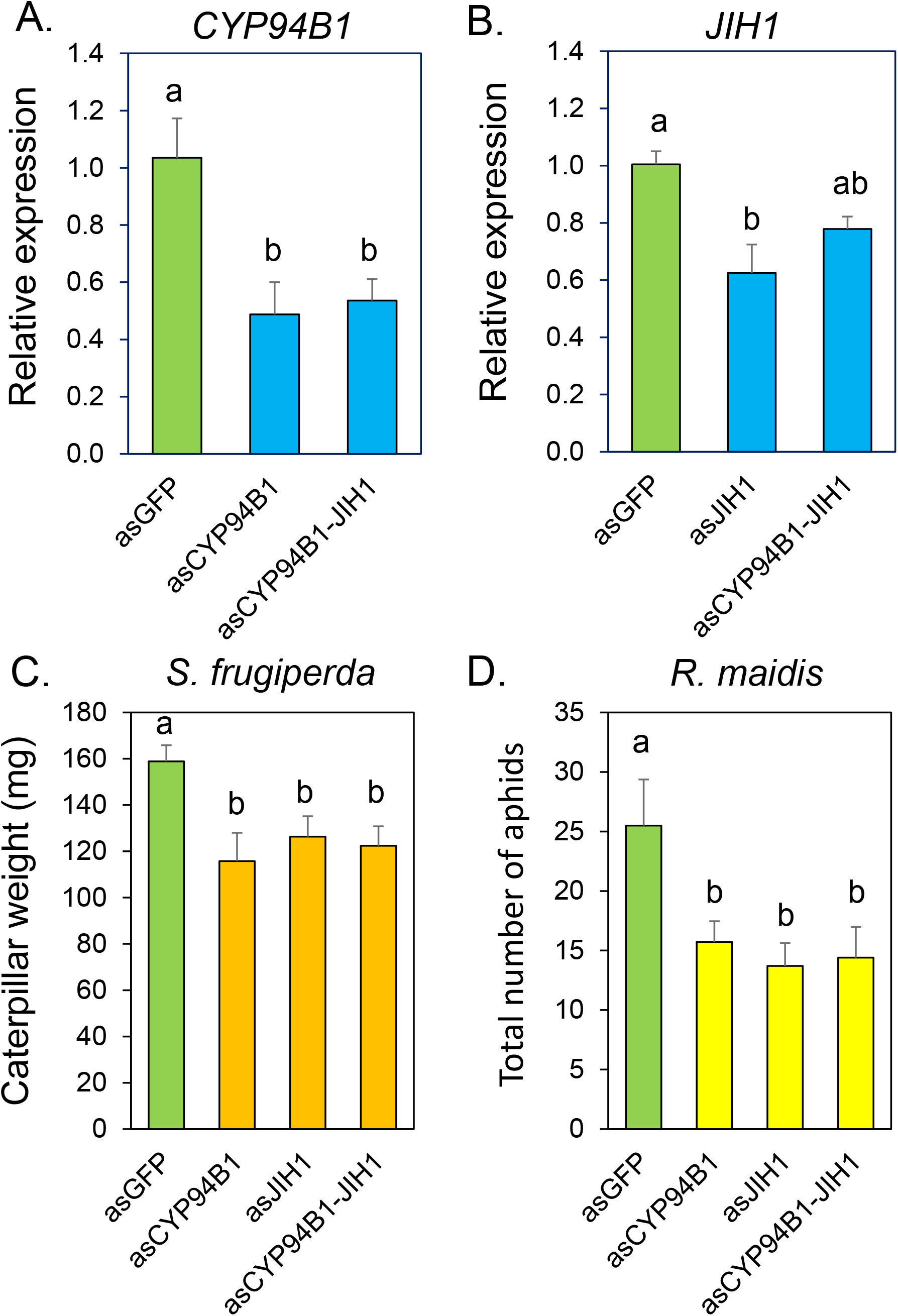
Simultaneous VIGS of *JIH1* and *CYP94B1* and its effects on insect growth. Maize inbred line P39 was rub-inoculated with SCMV carrying constructs targeting *JIH1, CYP94B1*, or both genes, with GFP as a control, in the antisense orientation. (A) *JIH1* expression and(B) *CYP94B1* expression, mean +/- s.e. of N = 5; (C) *Spodoptera frugiperda* mass after 1 week, mean +/- s.e. of N = 10, and (D) number of *Rhopalosiphum maidis* after 1 week. Mean +/- s.e. of N = 10-12; different letters indicate significant differences, P < 0.05, ANOVA followed by Tukey’s HSD test.

Together, our results show that, in addition to functioning as a gene overexpression vector (Mei et al., 2019; Mohr, 2019; Beernink et al., 2021; Chung et al., 2021), SCMV is effective for maize gene expression silencing. SCMV-induced gene expression silencing will be a useful tool for studying the *in vivo* function of maize genes without having to go through the lengthy and expensive process of making transgenic plants. Similar to what has been observed in Arabidopsis, rice, and *N. attenuata*, silencing the expression of *ZmCYP94B1* and *ZmJIH1* by SCMV-mediated VIGS indicates a role for the gene products in regulating maize defense. However, further work needs to be done to determine the more precise functions of these enzymes in attenuating JA-regulated defense responses.

## Acknowledgements

This work was supported by agreement HR0011-17-2-0053 from the Defense Advanced Research Projects Agency (DARPA) Insect Allies Program with the Boyce Thompson Institute, and United States Department of Agriculture (USDA) – National Institute of Food and Agriculture award 2021-67014-342357 and US National Science Foundation (NSF) award IOS-1339237 to GJ. HS was supported by the NSF DBI-2021795 and the USDA Hatch Grant TEX0-1-6584.

## Author Contributions

SHC, SAW, and GJ conceived of the project, SHC and SZ conducted experiments and interpreted data, HS provided essential research materials and interpreted data, SHC and GJ wrote the manuscript.

## Conflict of Interest Statement

The authors do not have conflicts of interest related to this project.

## Notes

### Competing Interest Statement

The authors have declared no competing interest.

## References

Ahmed F, Senthil-Kumar M, Dai X, Ramu VS, Lee S, Mysore KS, Zhao PX (2020) pssRNAit: A web server for designing effective and specific plant siRNAs with genome-wide Off-target assessment. Plant Physiol 184:65–81

Beernink BM, Holan KL, Lappe RR, Whitham SA (2021) Direct agroinoculation of maize seedlings by injection with recombinant foxtail mosaic virus and sugarcane mosaic virus infectious clones. J Vis Exp 168:e62277

Borrego EJ, Kolomiets MV (2016) Synthesis and functions of jasmonates in maize. Plants (Basel) 5:doi: 10.3390/plants5040041

Burch-Smith TM, Anderson JC, Martin GB, Dinesh-Kumar SP (2004) Applications and advantages of virus-induced gene silencing for gene function studies in plants. Plant J 39:734–746

Caarls L, Elberse J, Awwanah M, Ludwig NR, de Vries M, Zeilmaker T, Van Wees SCM, Schuurink RC, Van den Ackerveken G (2017) Arabidopsis JASMONATE-INDUCED OXYGENASES down-regulate plant immunity by hydroxylation and inactivation of the hormone jasmonic acid. Proc Natl Acad Sci U S A 114:6388–6393

Chen W, Shakir S, Bigham M, Richter A, Fei Z, Jander G (2019) Genome sequence of the corn leaf aphid (*Rhopalosiphum maidis* Fitch). Gigascience 8

Chuang WP, Herde M, Ray S, Castano-Duque L, Howe GA, Luthe DS (2014) Caterpillar attack triggers accumulation of the toxic maize protein RIP2. New Phytol 201:928–939

Chung SH, Bigham M, Lappe RR, Chan B, Nagalakshmi U, Whitham SA, Dinesh-Kumar SP, Jander G (2021) A sugarcane mosaic virus vector for rapid in planta screening of proteins that inhibit the growth of insect herbivores. Plant Biotechnol J 19:1713–1724

Chung SH, Jander G (2022) Inhibition of *Rhopalosiphum maidis* (corn leaf aphid) growth on maize by virus-induced gene silencing with sugarcane mosaic virus. Methods Mol Biol 2360:139–153

Cordero MJ, Raventos D, San Segundo B (1994) Expression of a maize proteinase inhibitor gene is induced in response to wounding and fungal infection: systemic wound-response of a monocot gene. Plant J 6:141–150

Ding XS, Mannas SW, Bishop BA, Rao X, Lecoultre M, Kwon S, Nelson RS (2018) An improved *Brome mosaic virus* silencing vector: Greater insert stability and more extensive VIGS. Plant Physiol 176:496–510

Felsenstein J (1985) Confidence Limits on Phylogenies: An Approach Using the Bootstrap. Evolution 39:783–791

Hayward A, Padmanabhan M, Dinesh-Kumar SP (2011) Virus-induced gene silencing in nicotiana benthamiana and other plant species. Methods Mol Biol 678:55–63

Hazman M, Suhnel M, Schafer S, Zumsteg J, Lesot A, Beltran F, Marquis V, Herrgott L, Miesch L, Riemann M, Heitz T (2019) Characterization of jasmonoyl-isoleucine (JA-Ile) hormonal catabolic pathways in rice upon wounding and salt stress. Rice (N Y) 12:45

Howe GA, Jander G (2008) Plant immunity to insect herbivores. Ann Rev Plant Biol 59:41–66

Jarugula S, Willie K, Stewart LR (2018) *Barley stripe mosaic virus* (BSMV) as a virus-induced gene silencing vector in maize seedlings. Virus Genes 54:616–620

Kant R, Dasgupta I (2019) Gene silencing approaches through virus-based vectors: speeding up functional genomics in monocots. Plant Mol Biol 100:3–18

Kitaoka N, Matsubara T, Sato M, Takahashi K, Wakuta S, Kawaide H, Matsui H, Nabeta K, Matsuura H (2011) Arabidopsis CYP94B3 encodes jasmonyl-L-isoleucine 12-hydroxylase, a key enzyme in the oxidative catabolism of jasmonate. Plant Cell Physiol 52:1757–1765

Koo AJ, Cooke TF, Howe GA (2011) Cytochrome P450 CYP94B3 mediates catabolism and inactivation of the plant hormone jasmonoyl-L-isoleucine. Proc Natl Acad Sci U S A 108:9298–9303

Koo AJ, Howe GA (2009) The wound hormone jasmonate. Phytochemistry 70:1571–1580

Koo AJ, Thireault C, Zemelis S, Poudel AN, Zhang T, Kitaoka N, Brandizzi F, Matsuura H, Howe GA (2014) Endoplasmic reticulum-associated inactivation of the hormone jasmonoyl-L-isoleucine by multiple members of the cytochrome P450 94 family in Arabidopsis. J Biol Chem 289:29728–29738

Kumagai MH, Donson J, della-Cioppa G, Harvey D, Hanley K, Grill LK (1995) Cytoplasmic inhibition of carotenoid biosynthesis with virus-derived RNA. Proc Natl Acad Sci U S A 92:1679–1683

Livak KJ, Schmittgen TD (2001) Analysis of relative gene expression data using real-time quantitative PCR and the 2(-Delta Delta C(T)) method. Methods 25:402–408

Lunde C, Kimberlin A, Leiboff S, Koo AJ, Hake S (2019) *Tasselseed5* overexpresses a wound-inducible enzyme, ZmCYP94B1, that affects jasmonate catabolism, sex determination, and plant architecture in maize. Commun Biol 2:114

Marquis V, Smirnova E, Graindorge S, Delcros P, Villette C, Zumsteg J, Heintz D, Heitz T (2021) Broad-spectrum stress tolerance conferred by suppressing jasmonate signaling attenuation in Arabidopsis JASMONIC ACID OXIDASE mutants. Plant J

Marquis V, Smirnova E, Poirier L, Zumsteg J, Schweizer F, Reymond P, Heitz T (2020) Stress- and pathway-specific impacts of impaired jasmonoyl-isoleucine (JA-Ile) catabolism on defense signalling and biotic stress resistance. Plant Cell Environ 43:1558–1570

Mei Y, Liu G, Zhang C, Hill JH, Whitham SA (2019) A *Sugarcane mosaic virus* vector for gene expression in maize. Plant Direct 3:e00158

Mei Y, Zhang C, Kernodle BM, Hill JH, Whitham SA (2016) A *Foxtail mosaic virus* vector for virus-induced gene silencing in maize. Plant Physiol 171:760–772

Mlotshwa S, Xu J, Willie K, Khatri N, Marty D, Stewart LR (2020) Engineering *Maize rayado fino virus* for virus-induced gene silencing. Plant Direct 4:e00224

Mohr I (2019) Examination of cucumber mosaic virus and sugarcane mosaic virus as VIGS and VOX vectors in Zea mays. MS. University of California, Davis, Davis, CA

Quilis J, Lopez-Garcia B, Meynard D, Guiderdoni E, San Segundo B (2014) Inducible expression of a fusion gene encoding two proteinase inhibitors leads to insect and pathogen resistance in transgenic rice. Plant Biotechnol J 12:367–377

Shivaji R, Camas A, Ankala A, Engelberth J, Tumlinson JH, Williams WP, Wilkinson JR, Luthe DS (2010) Plants on constant alert: elevated levels of jasmonic acid and jasmonate-induced transcripts in caterpillar-resistant maize. J Chem Ecol 36:179–191

Smirnova E, Marquis V, Poirier L, Aubert Y, Zumsteg J, Menard R, Miesch L, Heitz T (2017) Jasmonic Acid Oxidase 2 Hydroxylates Jasmonic Acid and Represses Basal Defense and Resistance Responses against Botrytis cinerea Infection. Mol Plant 10:1159–1173

Staswick PE, Tiryaki I, Rowe ML (2002) Jasmonate response locus *JAR1* and several related Arabidopsis genes encode enzymes of the firefly luciferase superfamily that show activity on jasmonic, salicylic, and indole-3-acetic acids in an assay for adenylation. Plant Cell 14:1405–1415

Tamayo MC, Rufat M, Bravo JM, San Segundo B (2000) Accumulation of a maize proteinase inhibitor in response to wounding and insect feeding, and characterization of its activity toward digestive proteinases of Spodoptera littoralis larvae. Planta 211:62–71

Tamura K, Stecher G, Kumar S (2021) MEGA11: Molecular Evolutionary Genetics Analysis Version 11. Mol Biol Evol 38:3022–3027

Tang J, Yang D, Wu J, Chen S, Wang L (2020) Silencing JA hydroxylases in *Nicotiana attenuata* enhances jasmonic acid-isoleucine-mediated defenses against *Spodoptera litura*. Plant Divers 42:111–119

Tzin V, Hojo Y, Strickler SR, Bartsch LJ, Archer CM, Ahern KR, Zhou SQ, Christensen SA, Galis I, Mueller LA, Jander G (2017) Rapid defense responses in maize leaves induced by *Spodoptera exigua* caterpillar feeding. J Exp Bot 68:4709–4723

Vila L, Quilis J, Meynard D, Breitler JC, Marfa V, Murillo I, Vassal JM, Messeguer J, Guiderdoni E, San Segundo B (2005) Expression of the maize proteinase inhibitor (mpi) gene in rice plants enhances resistance against the striped stem borer (Chilo suppressalis): effects on larval growth and insect gut proteinases. Plant Biotechnol J 3:187–202

Wang R, Yang X, Wang N, Liu X, Nelson RS, Li W, Fan Z, Zhou T (2016) An efficient virus-induced gene silencing vector for maize functional genomics research. Plant J 86:102–115

Whelan S, Goldman N (2001) A general empirical model of protein evolution derived from multiple protein families using a maximum-likelihood approach. Mol Biol Evol 18:691–699

Woldemariam MG, Galis I, Baldwin IT (2014) Jasmonoyl-l-isoleucine hydrolase 1 (JIH1) contributes to a termination of jasmonate signaling in N. attenuata. Plant Signal Behav 9

Woldemariam MG, Onkokesung N, Baldwin IT, Galis I (2012) Jasmonoyl-L-isoleucine hydrolase 1 (JIH1) regulates jasmonoyl-L-isoleucine levels and attenuates plant defenses against herbivores. Plant J 72:758–767

Zhang J, Yu D, Zhang Y, Liu K, Xu K, Zhang F, Wang J, Tan G, Nie X, Ji Q, Zhao L, Li C (2017) Vacuum and co-cultivation agroinfiltration of (germinated) seeds results in tobacco rattle virus (TRV) nediated whole-plant virus-induced gene silencing (VIGS) in wheat and maize. Front Plant Sci 8:393

